# Dynamic of the transcriptomic landscape of OsHV-1 replication in haemocytes of Pacific oyster

**DOI:** 10.1101/2025.07.11.664376

**Authors:** Aurélie Dotto-Maurel, Jérémy Le Luyer, Nicole Faury, Lionel Degremont, Margot Tragin, Tristan Renault, Benjamin Morga, Germain Chevignon

## Abstract

Since the 1990s, the Pacific oyster (*Magallana gigas*) has experienced repeated mortality events associated with Ostreid herpesvirus 1 (OsHV-1). Although the virus has been genomically characterised, its replication cycle and its interactions with the oyster immune system are still not well understood. In particular, little is known about the dynamics of OsHV-1 gene expression and the immune responses of haemocytes from oysters with varying susceptibility to the virus. While some studies have focused on the expression of specific viral and host genes on whole oysters, none have provided a comprehensive analysis of genomes-wide expression across multiple post-infection time points in haemocytes.

The lack of oyster cell lines makes studying virus-host interactions in vitro challenging. However, haemocytes, the key immune cells circulating in hemolymph, can be maintained *in vitro* in the short term and represent a relevant model for analyzing infection dynamics. In this study, haemocytes from two *M. gigas* families, one highly susceptible and one less susceptible to OsHV-1, were infected *in vitro*. We tracked the viral and host transcriptomes over a 24-hour period post-infection using high-throughput dual transcriptomics.

Our results provide a detailed overview of the OsHV-1 transcriptomic landscape in haemocytes from high and low susceptible *M. gigas* over time. In addition, WGCNA analysis of host genes expression provided insights into the haemocytes response to infection, and highlighted family-specific immune responses. This comprehensive transcriptomic study is the first to describe virus-host interactions across multiple stages of infection in haemocytes from Pacific oysters showing contrasted survival when exposed to OsHV-1.

**IMPORTANCE:** This study provides valuable insights into the interaction between *M. gigas* and OsHV-1 by analyzing viral expression and host immune response at the cellular level. By focusing on haemocytes, the key immune cells in Pacific oysters, the results reveal a link between host genotype and viral transcriptomic activity, providing new perspectives on molecular basis of natural susceptibility levels to OsHV-1 infection depending of the genetic background. Overall, our findings deepen the understanding of OsHV-1 gene expression dynamics and antiviral defense mechanisms in key species cultivated worldwide.

## INTRODUCTION

Herpesviruses are large, double-stranded DNA viruses known for their complex genome and biology (Silva et al., 2022). Their viral life cycle alternates between a lytic phase, which is characterized by active replication and spread, and a persistence phase during which the virus remains dormant and evades detection by the immune system. Research focused on the lytic phase of human herpesviruses, particularly Herpes Simplex Virus 1 (HSV-1) (Everett, 2014; Knipe et al., 2013) showed that this phase involves a tightly regulated transcriptional cascade comprising three gene classes: (i) immediate-early genes, which initiate virion replication by activating early gene transcription; (ii) early genes, which code proteins necessary for viral DNA replication; and (iii) late genes, which produce virion structural components such as capsid proteins and glycoproteins (Everett, 2014; P. C. Jones & Roizman, 1979; Knipe et al., 2013; Packard & Dembowski, 2021; Wofford et al., 2020)

While much attention has been given to herpesviruses that infect humans, these represent only eight out of the 137 known herpesvirus species. The *Herpesviridae* family has a broad host range and infects mammals, birds, fish, amphibians and marine molluscs (Davison, 2002, 2010; Dotto-Maurel et al., 2025). In particular, since the 1990s, Ostreid herpesvirus 1 (OsHV-1) has emerged as a major pathogen in aquaculture, causing recurrent mass mortality events in Pacific oysters and resulting in significant economic losses. This herpesvirus belonging to the family *Malacoherpesviridae* is the only species in the genus *Ostreavirus* (Davison et al., 2005; Renault, 2024). To date less than 100 genomes are available in public database, with size ranging from 199 kb to 207 kb, and containing 123 to 128 unique Open Reading Frames (ORF). While 55% of the predicted genes have no assigned function yet, the remaining genes are associated with processes such as secretion, membrane interaction, DNA replication, and apoptosis inhibition (Burioli et al., 2017; Davison et al., 2005; Xia et al., 2015).

The life cycle of OsHV-1 is not fully understood and characterized. However, it is thought to follow the general life cycle of other herpesviruses comprising both lytic and persistent phases. Indeed, when the seawater temperature reaches and remains above 16°C, this virus replicates actively and induces high mortality in juvenile oysters within two weeks in an intertidal environment. This occurs even more quickly under laboratory conditions, where the oysters are constantly immersed (Dégremont, 2013; Petton et al., 2013; Renault et al., 1994, 2011; Segarra, Mauduit, et al., 2014). However, when the temperature of seawater falls below 13°C, no mortality is observed and OsHV-1 becomes largely undetectable suggesting that it enters a persistent stage (Dégremont et al., 2013; Degremont & Benabdelmouna, 2014; Petton et al., 2013; Renault et al., 2014).

During the lytic phase, it has been suggested that OsHV-1 enters the Pacific oyster *via* the haemolymphatic system, and then infect various organs throughout the hemolymph (Schikorski, Faury, et al., 2011; Segarra et al., 2016). *In situ* hybridization studies and DNA detection by qPCR have shown that OsHV-1 is able to replicate in several oyster tissues, in particular in the gills, mantle, labial palps, digestive gland, gonad, heart, adductor muscle and ganglia (Lipart & Renault, 2002; Martenot et al., 2016; Segarra et al., 2016). Techniques such as reverse transcription real-time PCR (RT-qPCR), microarray and RNAseq have been used to study viral and oyster gene expressions (de Lorgeril et al., 2018; Jouaux et al., 2013; Morga et al., 2017; Renault et al., 2011; Segarra, Faury, et al., 2014; Segarra, Mauduit, et al., 2014). Those approaches have shown that OsHV-1 gene expressions occur at an early stage in the infection and have enabled the classification of some OsHV-1 genes as either immediate-early or early (Morga et al., 2017; Segarra, Faury, et al., 2014). While these findings have deepened our understanding of OsHV-1 gene expression dynamics during infection, they also raise important questions about the molecular mechanisms by which oysters detect and respond to viral invasion.

Antiviral immunity in mollusks is still not well understood. However, Pacific oysters have a sophisticated and well-coordinated defense system that can detect nonspecific nucleic acids and activate an antiviral response that inhibits subsequent OsHV-1 infection (Green et al., 2015; Segarra, Mauduit, et al., 2014). Moreover, it has been shown that OsHV-1 infection leads to increase expression of various genes associated with an interferon-like pathway, including viral recognition receptors, signal transducers, transcription factors, and antiviral effectors (Green et al., 2015; Rosani et al., 2015; Wang et al., 2018). However, while these molecular findings highlight the existence of a complex antiviral response in oysters, they do not fully explain the variability observed in disease outcomes between host with different genetic background.

Indeed, susceptibility to OsHV-1 infection varies significantly among Pacific oyster families, suggesting that genetic background plays a crucial role in determining the effectiveness of the host immune response. While some families are highly susceptible to OsHV-1 infection and experience high mortality rates, others show much higher survival rates. Selective breeding has enabled the production of oyster families that demonstrate contrasted susceptibilities to OsHV-1 infection (Dégremont, 2011; Dégremont et al., 2015). Although low susceptible Pacific oysters are not completely resistant to infection and still carry the virus, they are better at controlling viral replication than more susceptible ones (Dégremont, 2011; Segarra, Mauduit, et al., 2014). Indeed, transcriptomic analysis showed that low susceptible Pacific oysters exhibit a faster and more effective immune response than more susceptible ones. However, these analyses were conducted on whole-oyster tissue pools, which limits the ability to assess tissue-specific resolution of the immune response (de Lorgeril et al., 2018).

Currently, no cell line system is available to study OsHV-1 replication *in vitro*, making transcriptomic analysis of the lytic phase challenging. However, haemocytes, the circulating cells present in Pacific oyster haemolymph, can be maintained *in vitro* several hours and can be infected by OsHV-1 to study the lytic phase within simplified model (Morga et al., 2017). Although haemocytes play a crucial role in the immune response of *M. gigas*, only few studies have investigated their response to OsHV-1 infection.

Our study aims to provide a detailed overview of the OsHV-1 transcriptomic landscape in primary culture of haemocytes from low susceptible (LS) and high susceptible (HS) Pacific oysters over a 24-hour infection period. RNA from haemocytes was extracted and sequenced using short-read technology to analyse viral expression and host response. This dual transcriptional analysis represents the first characterization of host-virus interactions in two phenotypically distinct oyster families in terms of viral infection susceptibility over multiple infection times under *in vitro* conditions.

## MATERIALS & METHODS

### 1. Pacific oysters production

One batch of *M. gigas* selected for its higher susceptibility (HS) to OsHV-1 infection, and one batch selected for its lower susceptibility (LS) were conditioned at the Ifremer hatchery in La Tremblade in December 2014. The two batches correspond to the control and selected groups produced in the fourth generation of the line B described in a previous study (Dégremont et al., 2015). The broodstock were maintained in 240 L raceways with continuous supply of filtered seawater treated with UV and were richly fed with phytoplankton (*Isochrysis galbana, Tetraselmis suecica,* and *Skeletonema costatum*) to promote gametogenesis. Before reproduction, twelve oysters per family were tested for the detection of *Vibrio aestuarianus* and OsHV-1 using standard qPCR protocols (Pepin et al., 2008; Saulnier et al., 2009).

Spawning occurred on March 11, 2015. For each family, 25 to 30 oysters were placed in a 5 L beaker and induced to spawn using heat shock by changing the seawater temperature from 12°C to 28°C. Once spawning began, the seawater temperature was maintained at 25°C. Gametes were sieved through a 100 µm sieve to remove larger tissue debris and then a 20 µm sieve to remove small tissue/sperm debris. Fertilized and unfertilized eggs were retained on the 20 µm sieve. The embryos were then transferred to a 30 L tank with UV-filtered seawater for which temperature was maintained at 25°C. Seawater was changed three times per week. Larvae were fed daily with *I. galbana*, *T. suecica* and *S. costatum* was added to the feed when larvae size exceeded 140 µm. Two weeks after fertilization, pediveliger larvae were transferred to sieves in 120 L tanks, and raised under standard conditions in our controlled facilities (UV-treated seawater).

### 2. Haemolymph collection

At 8 month old, and for each family, the haemolymph from 450 oysters, approximately 0.5 to 1 mL per oyster, both families was collected from the adductor muscle sinus using a 1 mL syringe equipped with a 20G AGANI ™ needle (0.9 x 40 mm). To remove haemolymph debris, samples were filtered through a 60 µm nylon mesh and kept on ice to limit cell aggregation. Then, haemolymph samples were pooled within family, and haemocytes were counted using a Malassez cell. The haemocytes concentration was adjusted to 1.10^6^ cells per mL using filtered 0.22 µm artificial seawater (AWS) (Figure 1).

**Figure 1:**
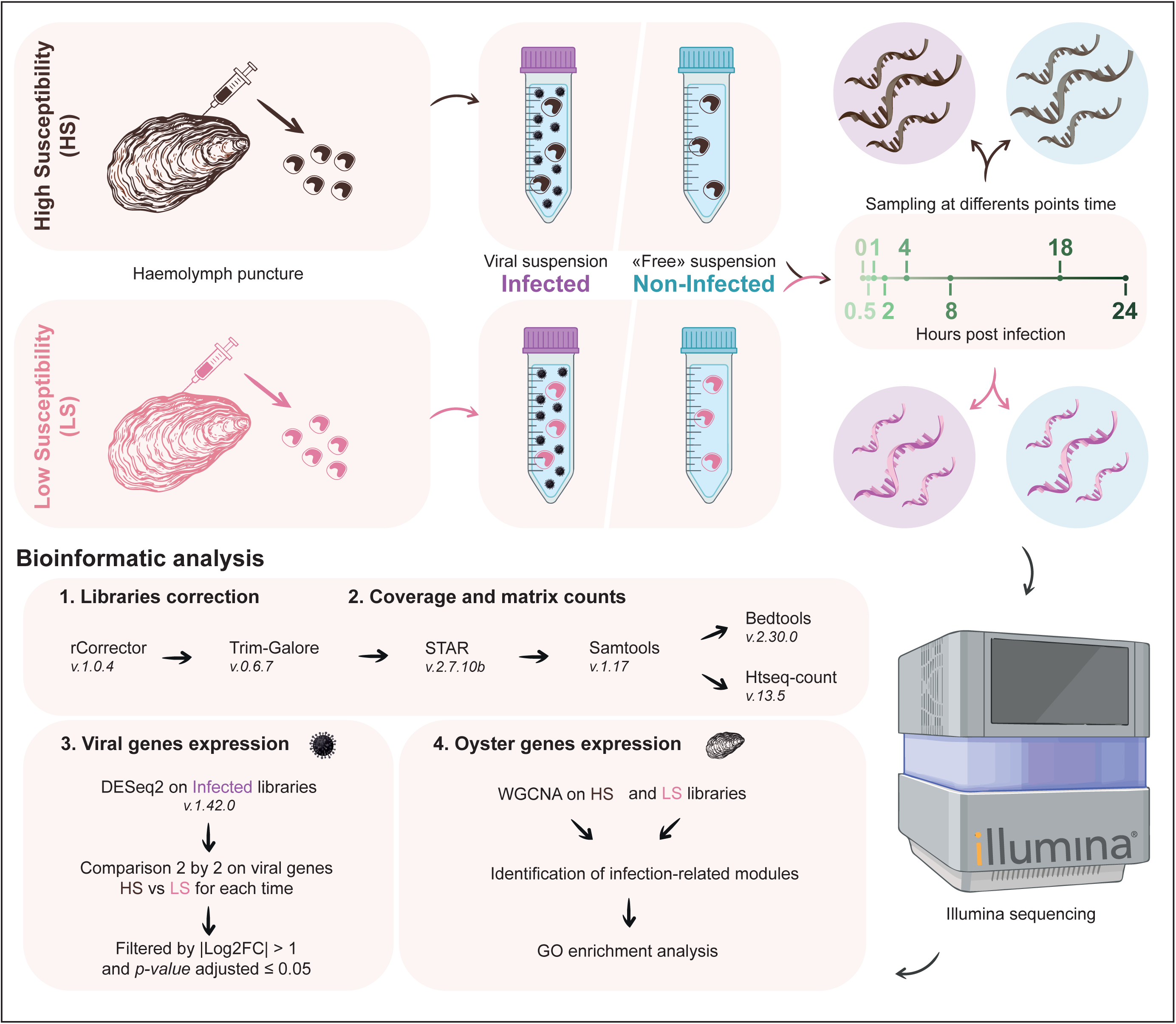
Experimental design and bioinformatic pipeline implemented in this study.

### 3. Viral suspension

Viral suspension was prepared from infected oysters following the method outlined by Schikorski et al. (Schikorski, Renault, et al., 2011). To produce controls, the protocol described above was applied to oysters that were tested negative for the detection of OsHV-1 DNA by real time PCR to generate a virus-free suspension.

### 4. *In vitro* infection

For each family, the pool of haemocytes were separated in 3 replicates for each time and each treatment (i.e. infected and non-infected). The infected (Inf) haemocytes (1.10^6^ cells/mL, 5 mL) were incubated with viral suspension (2.5 mL, 10^5^ copies of OsHV-1/µL) under gentle agitation at 19°C and sampled after 0 min, 30 min, 1 hr, 2 hrs, 4 hrs, 8 hrs, 18 hrs, and 24 hrs of *in vitro* virus exposure (n=3 for each condition except at 30 min where n=2). Similarly, the non-infected (nI) haemocytes (1.10^6^ cells/mL, 5 mL) were incubated using the virus-free suspension (2.5 mL) and sample at the same time steps (Figure 1).

To prevent fungal and bacterial growth in all samples, 350µl of antibiotics was added, including 4 mg/mL of streptomycin, 11.6 mg/mL of penicillin, 5.1 mg/mL of neomycin, 3.3 mg/mL of erythromycin, and 0.1 µL/mL of nystatin.

### 5. Haemocytes DNA extraction and OsHV-1 DNA quantification

At each sample time, 7.5 ml of haemolymph was centrifuged for 10 min at 1 500 g, the pellet was then DNA extracted using the QiAmp Tissue Mini Kit (QiAgen) following the manufacturer’s protocol.

To assess the viral genome copy number, a real-time quantitative PCR (qPCR) using a Mx3005P Thermocycler sequence detector (Agilent), was performed on extracted DNA.

Amplification reactions were performed in a total volume of 20 μL. Each well contained 5 μL DNA from sea water or 5ng of total DNA from oyster mantle, 10 μL of Brilliant III Ultra-Fast SYBR®Green PCR Master Mix (Agilent), 2 μL of each primer OsHVDP For (forward) 5’-ATTGATGATGTGGATAATCTGTG-3’ and OsHVDP Rev (reverse) 5’-GGTAAATACCATTGGTCTTGTTCC-3’ (Webb et al., 2007) at the final concentration of 550 nM each, and 1 μL of distilled water. Real time PCR cycling conditions were as follows: 3 min at 95°C followed by 40 cycles of amplification at 95°C for 5 s and 60°C for 20 s.

### 6. RNA extraction and sequencing

Similar to DNA extraction, 7.5 ml of haemolymph was centrifuged for 10 min at 1 500 g and the Total RNA from the pellet was extracted using TRIZOL® Reagent™ (Ambion®) following the manufacturer’s recommendations. RNA was then treated with Turbo™ DNase (Ambion®) to remove any remaining DNA. After DNase treatment, a second RNA purification using TRIZOL was performed. The quality and quantity of the RNA were checked using Nanodrop. First-strand cDNA synthesis was carried out using the SuperScript® III First-Strand Synthesis System from Invitrogen with 500 ng of RNA. A No RT (No Reverse Transcription) control was performed after RNA extraction and was verified by qPCR to confirm the absence of oyster and/or virus genomic DNA using EF1α primers (targeting the elongation factor 1 alpha, Forward: 5’-GTCGCTCACAGAAGCTGTACC-3’, Reverse: 5’-CCAGGGTGGTTCAAGATGAT-3’) and OsHVDP For/OSHVDP Rev primers (Targeting the ORF 100 coding for DNA polymerase: OsHVDP For (forward) 5’-ATTGATGATGTGGATAATCTGTG-3’; OsHVDP Rev (reverse) 5’-GGTAAATACCATTGGTCTTGTTCC-3’ (Webb et al., 2007)).

Sequencing of the 94 libraries was performed by Eurofins Genomics using strand-specific cDNA sequencing on Illumina HiSeq 2500/Illumina MiSeq.

### 7. Data Analysis

All scripts described in this part are detailed in Gitlab (https://gitlab.ifremer.fr/asim/haemocytes-dualrna-seq-analysis.git) and illustrated in Figure 1.

#### Annotation of the OsHV-1 genome

The experiment described in this study was carried out in 2015. For more accurate data analysis, we used an OsHV-1 reference as close as possible to the haplotype used at that time (Raw data: SAMN18712119) (Morga et al., 2021), which was assembled from viral DNA extracted from infected oysters in the Bay of Marennes-Oléron in 2017 (Morga et al., 2021).

This genome was then annotated using pairwise alignment with the OsHV-1 µVar A genome (Burioli et al., 2017) using Geneious version 2025.0.2 (https://www.geneious.com). The annotation from the OsHV-1 µVar A genome was then transferred to the OsHV-1 genome used in this study. To refine the annotation, four ORF predictors were used: FragGeneScan (Rho et al., 2010), Vgas (K.-Y. Zhang et al., 2019), Prodigal (https://github.com/hyattpd/Prodigal) and GeneMark (https://exon.gatech.edu/GeneMark/gmhmme.cgi). Following prediction, ORFs that were added or modified relative to the OsHV-1 µVarA genome were labelled as ‘ORFS’ to indicate Supplementary ORF (Table S1).

The functions and domains of all ORFs were then predicted using Interproscan version 5.73-104.0 (P. Jones et al., 2014) with all database available and EggNogg mapper version 2.1.5 – 2.1.12 (Cantalapiedra et al., 2021) web tools interrogated on February 2025. In addition, DeepLoc version 2.1 (https://services.healthtech.dtu.dk/services/DeepLoc-2.1/) was used to predict the subcellular localizations of viral proteins (Table S1).

#### RNAseq reads mapping, reads count and descriptive statistics

The sequencing libraries were corrected using rCorrector (Song & Florea, 2015) and Trim Galore (https://github.com/FelixKrueger/TrimGalore) with default parameters to remove adapters introduced during sequencing and to filter out short and poor quality reads (Script 01-Correct_libraries in Gitlab). Reads were then aligned to the *M. gigas* mitochondrial genome (GenBank: GCF_963853765.1), the *M. gigas* nuclear genome (GenBank: NC_001276) and the OsHV-1 genome (Raw data: SAMN18712119) (Morga et al., 2021) simultaneously, using STAR (Dobin et al., 2013) with default parameters.

Forward and reverse count matrices were generated for each of the 94 datasets using HTSeq-count v.0.13.5 (Anders et al., 2015), with reads counted at the gene level using the –t exon, –i gene and –s yes or –s reverse options. All forward and reverse matrices were then combined to produce two complete count matrices covering all 94 datasets (Script 02-Alignments_Count_matrices). Outlier libraries were identified from the count matrix by plotting heatmap and scatterplots adapted from the iDEP 2.01 scripts (http://bioinformatics.sdstate.edu/idep/).

Four different factors were then evaluated: (1) “treatment” representing OsHV-1 infected or non-infected hemocytes, (2) “viral load” corresponding to the amount of OsHV-1 genome copy number per nanograms of DNA, (3) “time” which correspond to the 8 sample time points (0, 0.5, 1, 2, 4, 8, 18, 24 hours post infection (hpi)) and (4) “family” traducing contrasted phenotype of the Pacific oysters: high or low susceptible to OsHV-1 infection.

To assess the relative contribution of each of these factors to the gene expression levels, a distance-basedredundancy analysis (db-RDA) was performed. First, the “time” factor was centred and scaled (sc). The data were then normalized using DESeq2 ‘median of ratios’ method as implemented in R package (Love et al., 2014) with “∼treatment*sc_time” as the design formula. The normalized count matrix was then transformed using a variable-stabilizing transformation (vst), followed by principal coordinate analysis (PCoA). on the Euclidean distances A db-RDA was then performed using the retained PCoA factors as the response matrix and the variables “time”, “family” and “viral load” as the explanatory matrix. A total of six PCoA axes were retained based on the Gower dissimilarity index, explaining 66.86% of the total variance. Three partial db-RDAs and permutation test for constrained ordination (N = 1000 permutations) were performed to validate the effect of each factor while controlling for others.

All R analyses described in this section can be found in script 05-R_script_1_sequencing_analysis_raw on GitLab.

#### OsHV-1 gene expression analysis

To access OsHV-1 RNAseq coverage, the.sam files produced by STAR were processed using Samtools v.1.16.1 (Danecek et al., 2021) with parameters samtools view –f 128 and –F 16, –f 64 and –F 32, –f 144, –f 96, to select forward and reverse aligned reads and samtools merge –f to merge all forward and all reverse aligned reads. Then bed coverage files (i.e. coverage per base on the OsHV-1 genome) were generated using Bedtools v.2.30.0 (Quinlan & Hall, 2010) with bedtools genomecov with standard parameter (Script 03-Generate_bed_files in Gitlab). Reads per million (RPM) normalization was applied to all bed coverage files by dividing the coverage at each base by the total number of reads in the library and multiplying the result by 1×10^6^.

For OsHV-1 genome coverage evaluation over time and across families, replicate data were averaged for each base of the genome. Antisense and sense reads of the OsHV-1 ORFs were identified using the orientation of the alignments and the OsHV-1 genome annotation with a custom script. The ratio of antisense to sense reads was also calculated for each position and each time point.

To compare viral ORFs expression over time and across families, the forward count matrix of all genes (i.e. *M. gigas* and viral ORFs) was normalized using DESeq2 version 1.46.0 (Love et al., 2014), with a design formula that included “time”, “family” and their interaction “time + family + time:family”. Viral ORFs were extracted from normalized count matrix and further normalized by the length of the OsHV-1 ORFs per kilobases. We first evaluate the expression of new and modified ORFs, considering an ORFs as expressed when the normalized count was greater than 1 RPKM.

To analyze viral gene expression profiles over time, we applied a variance-stabilizing transformation (vst) on the DESeq2 object containing all virus and host genes and extracted ORF data from this matrix. We then computed the Euclidean distances for average-linkage hierarchical clustering, removed ORFs that were never expressed at any time points from the dataset and cut the resulting dendrogram into three clusters.

Finally, the expression of ORFs in LS and HS families was compared by performing statistical tests at each time point on the DESeq2 object. An ORF was considered differentially expressed (DE) if the absolute value of its log_2_ fold-change was greater than 1 and if its adjusted p-value was less than 0.05. A negative log_2_ fold-change indicated that the ORF was overexpressed in the LS family, whereas a positive log_2_ fold-change indicated overexpression in the HS family.

All R analyses described in this section can be found in script 06-R_script_2_OsHV-1_ORFs_analysis_raw on GitLab.

#### *M. gigas* gene expression analysis

To analyze Pacific oyster gene expression, genes were extracted from the previous vst matrix. Low variance genes (i.e. genes with variance less than 0.05) were removed from the dataset.

First, to analyze the response of Pacific oyster genes to the viral infection according to the time for each family, we performed a weighted correlation network analysis (WGCNA) using the R package WGCNA (Langfelder & Horvath, 2008) on all infected libraries. To do so, we created a design matrix including “LS.time”, “HS.time”, “Cluster1”, “Cluster 2” and “Cluster 3” corresponding to the eigenvalues of each of the three viral clusters per libraries, and “family”. The soft power threshold was set at 10, using the scale-free topology criterion, achieving a model fit of 0.78 and an average connectivity (k) of around 275. Genes modules were identified using the cutreeDynamic function, with a minimum of 50 genes per module.

Then, to analyze the response of oyster genes to the time for each family we performed a WGCNA analysis using the R package WGCNA version 1.73 (Langfelder & Horvath, 2008) on all control libraries. To do so, we created a design matrix including “time” and “family”. The soft power threshold was set at 9, using the scale-free topology criterion, achieving a model fit of 0.76 and an average connectivity of around 379. Genes modules were identified using the cutreeDynamic function, with a minimum of 50 genes per modules.

For each module of each WGCNA analysis, the module membership was defined and the correlation between the module eigengene value and condition traits was assessed. Modules that showed a significant correlation (i.e. P<0.005) with interested traits were retained and genes belonging to these modules were extracted.

To perform functional enrichment analysis, Gene Ontology (GO) terms were assigned to Pacific oyster genes. Functional annotation of the oyster genome was performed using EggNOG-mapper v2 (Cantalapiedra et al., 2021) for orthology-based inference, and BLASTp (Altschul, 2014) searches against the curated UniProt-SwissProt database (released January 2023, (Bateman et al., 2021)) for high-confidence protein annotations (treshold e-value < 10e-5). Finally, a single file combining all GO annotations was generated with the nrifyGOtabl.pl script of GO_MWU tool (Wright et al., 2015) (Script 04-GO_annotations in Gitlab).

Custom scripts were then used to extract the best hits, retrieve GO terms from the BLAST and EggNogg-mapper outputs. Functional enrichment analysis was performed on these genes using a rank-based GO approach with adaptive clustering, applying a Mann-Whitney U test for each independent module using the GO_MWU R package (Wright et al., 2015) with the go.obo database downloaded on January 2025. REviGO was used to reduce redundancy in GO terms (Supek et al., 2011). To visualize functional enrichment, the negative logarithm of the adjusted p-value and the ratio of significant genes on total genes were calculated for each GO.

All R analyses described in this section can be found in script 07-R_script_3oyster_genes_analysis_raw on GitLab.

## RESULTS

### 1. Sequencing summary

#### Data quality and outlier removal

Of the 94 libraries (i.e. 46 for the LS family and 48 for the HS family), the total number of reads per library ranged from 9 to 20 million, with an average of 13 million reads. Alignments to the Pacific oyster and virus genomes showed that 88% to 94% of the reads of all libraries aligned to the Pacific oyster genome (average: 93%). For the infected condition, 0.00022% to 0.31651% of the reads from the LS family libraries aligned to the OsHV-1 genome (average: 0.083%), whereas 0.00023% to 2.69% of the reads from the HS family libraries aligned to the OsHV-1 genome (average: 0.39%). Finally, between 5.95% and 11.97% of the reads did not align to either *M. gigas* or OsHV-1 genomes in the LS family (average: 7.02%), and between 6.06% and 8.96% of the reads did not align to either *M. gigas* or OsHV-1 genomes in the HS family (average: 7.0%) (Figure S1).

By plotting the top 5000 most variable genes of our dataset on a heatmap, we identified and removed 5 outlier libraries: for the LS family, the first replicates of the non-infected condition at 1 hpi and 4 hpi, and the third replicate at 18 hpi; for the HS family, the second replicate of the non-infected condition at 18 hpi and the first replicate of the infected condition at 18 hpi (Figure S2).

#### Oyster family and time post-infection drive the variability of the whole expression

The RDA plot illustrates the variance in gene expression between oyster families under two treatments: non-infected and infected, across time points (Figure 2). The first axis of the RDA (RDA1), accounting for 58.1% of the variance, shows a clear separation between the LS and the HS family libraries. The second principal axis (RDA2), explaining 34.58% of the variance, further differentiates samples over time. These results suggest that both factors “families” and “time” significantly influence the gene expression patterns. This is verified by the adjusted R^2^ of the partial RDA showing that 14.40% of the variance is explained by the “time”, 15.73% by the “families” and 8.91 % by the “viral load”.

**Figure 2:**
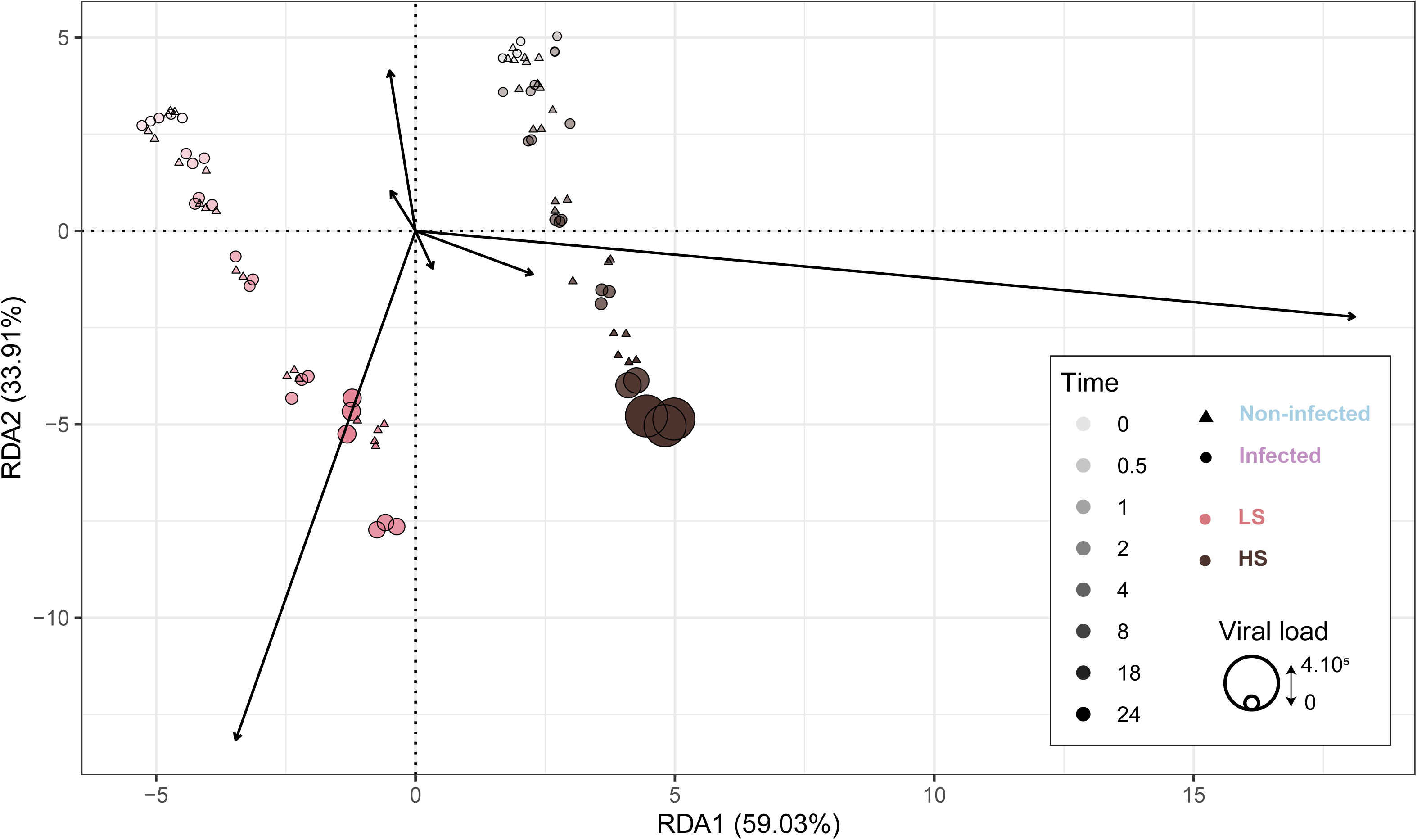
Redundancy Analysis (RDA) plot. The X-axis represents the first RDA component which explains 59.03% of the variance and the Y-axis represents the second RDA component, which explains 33.91%. Dot shape indicates treatment, with triangles for non-infected libraries and circles for infected libraries. Dot size corresponds to the viral load (i.e. OsHV-1 copy number per ng of extracted DNA). Color correspond to oyster family, with pink for LS family and darkbrown for HS family. The shade of the dot indicates the time after infection: light pink or brown dots correspond to early time points, while dark pink or brown dots correspond to later time points.

#### Viral genomes copy number and viral transcription levels are higher in the high susceptible family

Our qPCR results showed that the OsHV-1 copy number in DNA extracted from haemocytes of LS oysters increased slowly over time, reaching a maximum of 4.7.10^4^ cp/ng of DNA at 24 hpi. In contrast, OsHV-1 copy number increased rapidly to 4.3.10^5^ cp/ng of DNA in haemocytes from HS oysters (Figure 3.A).

**Figure 3:**
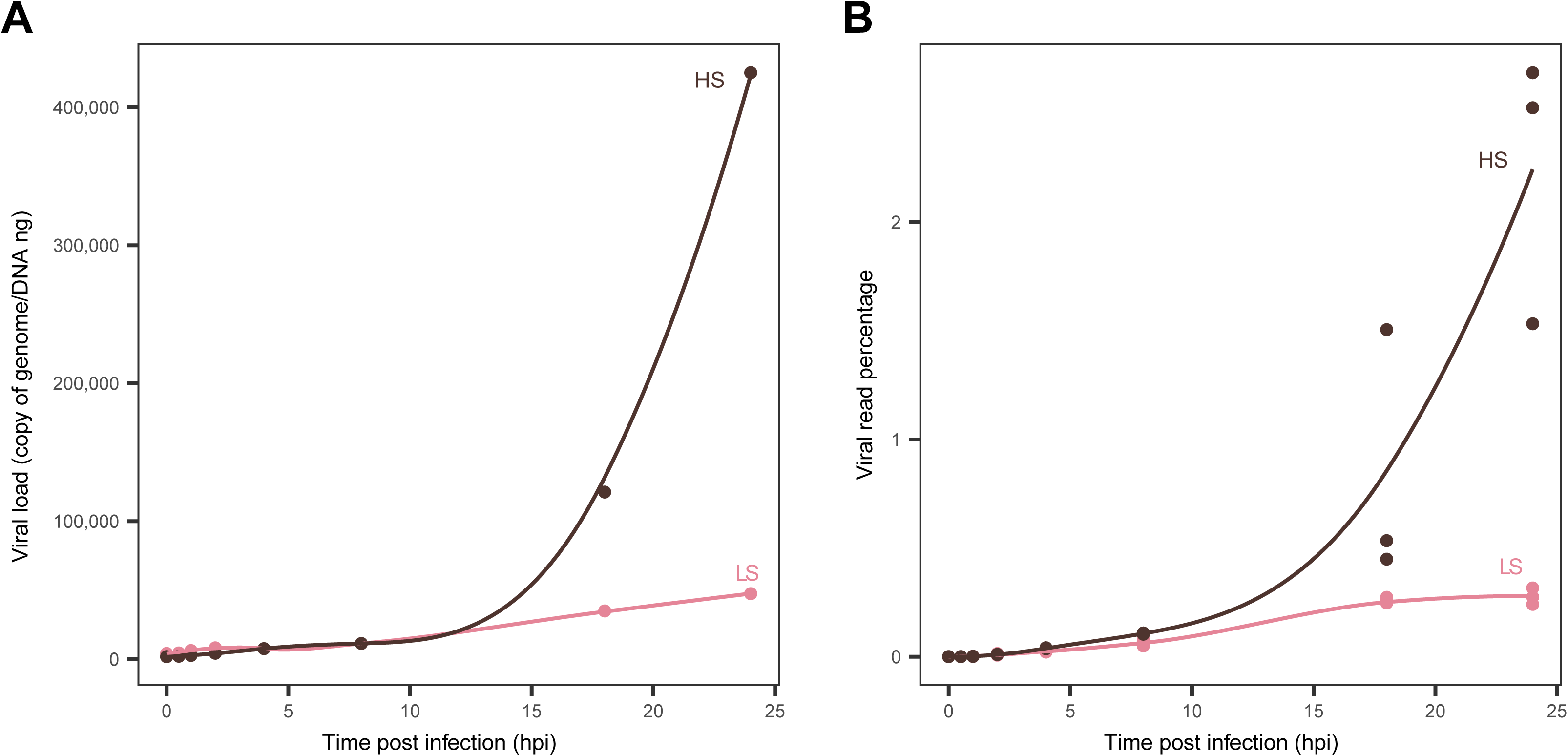
Viral DNA and RNA quantity over time. A. Viral load in copy of genome per ng of DNA over time. Each point represents a sample coloured by the haemocyte family they belong to (pink for LS family and darkbrown for HS family). The Y-axis represent viral DNA copies number per ng of DNA while the X-axis corresponds to time in hours post infection. B. Percentage of viral reads over time. Each point represents a cDNA library, colored by the family they belong to (pink for LS family and darkbrown for HS family). The Y-axis represents the percentage of OsHV-1 reads while the X-axis represents the time in hours post-infection.

A similar pattern appears in the percentage of viral reads from the RNAseq between the LS and the HS families. For the LS family, the percentage increased slowly to 0.3% at 24 hpi, whereas it increased quickly to a maximum of 2.6% at 24 hpi for the HS family (Figure 3.B).

### 2. OsHV-1 gene expression

#### Updated OsHV-1 gene prediction reveal new and modified ORFs

Our updated ORF prediction identified 144 ORFs, of which 5 were modified compared to the OsHV-1 µVar A genome (GenBank: KY242785) and 17 were newly predicted (Table S1). Of the new ORFs, three (ORFS4.2, ORFS105.3, ORFS116.1), exhibited expression levels below 1 RPKM in both families at all time points. ORFS32.1 was also lowly expressed in the LS family, but exceeded 1 RPKM at 18 hpi and 24 hpi in the HS family. The functions of these four ORFs are unknown, but predictions show they may be addressed to the endoplasmic reticulum (ORFS32.1) or in the nucleus (ORFS4.2, ORFS105.3 and ORF116.1) (Table S1).

The remaining 13 newly predicted ORFs and the 5 modified ORFs were expressed above 1 RPKM in both families. None of them have known functions, but they are predicted to localize in the cytoplasm or nucleus (ORFS32.2, ORFS33.1, ORFS119.2, ORFS50, ORFS69.1, ORFS72.1, ORFS105.4, ORFS4.1, ORFS43.3, ORFS115.2 and ORFS118.1); the extracellular region (ORFS121.1); and the membrane (ORFS62).

Of the five modified ORFs, only ORFS115 had a known function (an origin-binding replication protein; (Burioli et al., 2017)). The four others showed predicted features including a coiled-coil domain in ORFS17, a disordered region in ORFS32.3 and a signal peptide in ORFSIN1.2. Their predicted localizations were nuclear (ORFS17 and ORFS115), cytoplasmic (ORFS114), membrane (ORFS32.3) and extracellular (ORFSIN1.2) (Table S1).

#### OsHV-1 gene expression dynamics highlight differences between both Pacific oyster families

To investigate OsHV-1 gene expression dynamics, viral genome coverage and normalized counts were analyzed over time by focusing on the 46 OsHV-1 infected libraries.

Three distinct clusters of viral genes were identified based on their expression profiles (Table S1). Cluster 1 includes 9 genes, Cluster 2 includes 97 genes and Cluster 3 includes 38 genes (Figure 4). Cluster 1 genes are expressed immediately after infection with a median onset after 0 hpi. This cluster includes ORF27 coding dUTPase-like, predicted to localize in the cytoplasm; ORF45 and ORF107 coding proteins with disordered regions, predicted to localize in the nucleus or cytoplasm; ORF80 coding a transmembrane glycoprotein with a signal peptide, predicted to localize in the Golgi apparatus, ORF88 and ORF111 coding transmembrane glycoproteins and ORF82, ORF104 and ORF122 coding proteins of unknown function, predicted to localize in the nucleus (Figure 4, Table S1).

**Figure 4:**
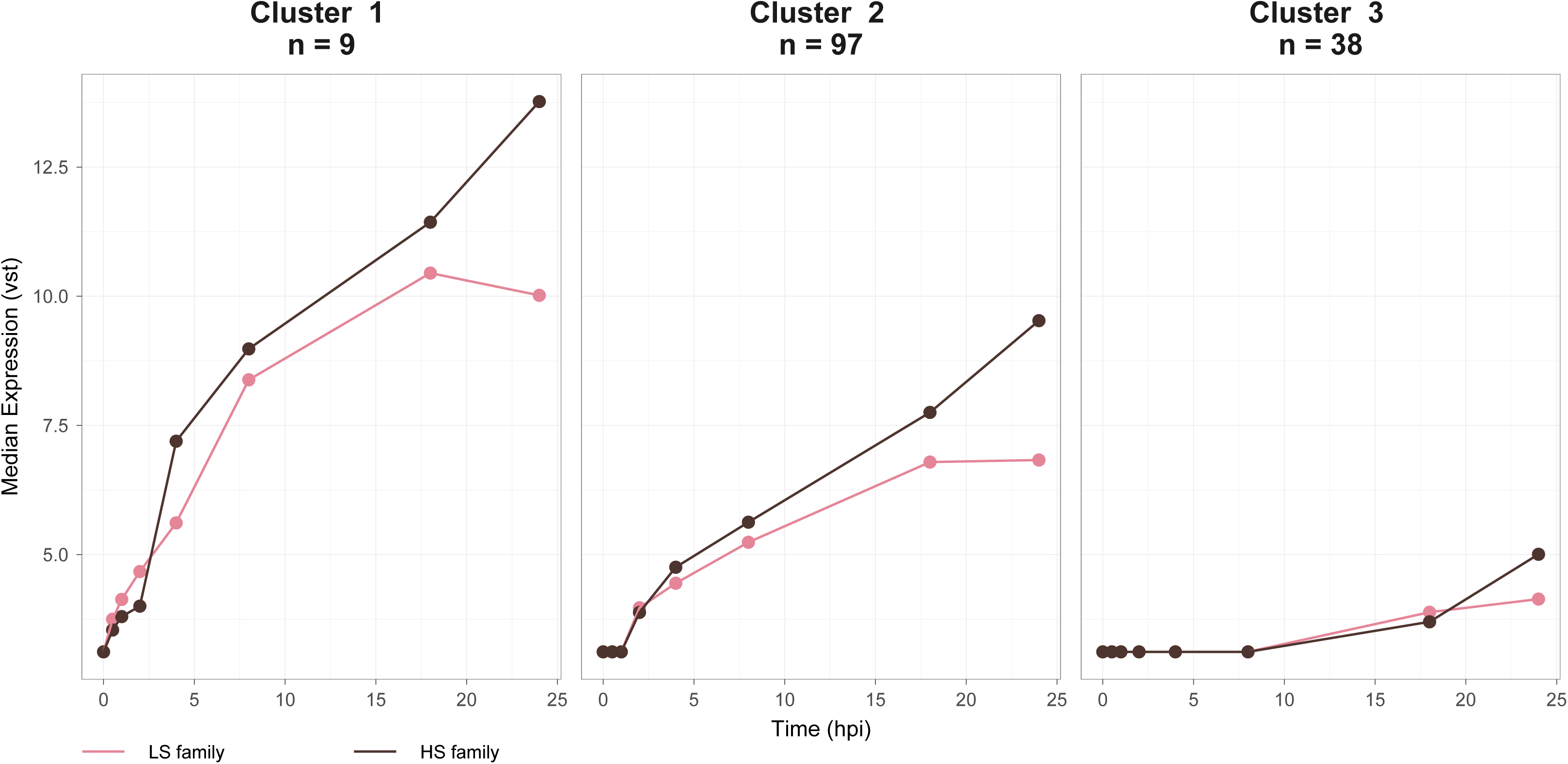
Clustering of viral genes based on their expression. Each panel shows one of the three clusters. The Y-axis shows the median expression, transformed using a variance-stabilizing transformation (VST), and the X-axis shows time in hours post-infection. Pink lines and dots represent the LS family, and dark brown lines and dots represent the HS family.

Cluster 2, the largest group, is characterized by early expression, with a median onset after 1 hpi. This cluster includes several key functional genes, such as ORF75, which encodes a dUTPase-like protein, four genes involved in the inhibition of apoptosis (ORF42, ORF87, ORF99 and ORF106), and eight out of the nine known genes involved in the DNA replication and coding for ribonucleotide reductase subunits (ORF20 and ORF51), DNA primases (ORF7, ORF49 and ORF66), DNA helicase (ORF67), DNA polymerase (ORF100) and a mitochondrion-localized P-loop containing nucleoside triphosphate hydrolase (ORF44) all predicted to localize in the cytoplasm and/or nucleus (Figure, 4, Table S1).

Cluster 3 genes are predominantly expressed at later stages of infection, with a median onset after 8 hpi in both families. Notably, this cluster includes 15 modified or newly annotated ORFs, as well as ORF109, which codes a putative DNA packaging protein (Figure 4, Table S1).

Expression dynamics of clusters 2 and 3 differ between host families. In the LS family, genes from clusters 2 and 3 show increasing expression up to 18 hpi, followed by a decline at 24 hpi. In contrast, expression in the HS family continues to rise until 24 hpi. Overall, viral gene expression levels are lower in the LS family than in the HS one (Figure 4).

By the end of infection, five ORFs (ORF13, ORF27, ORF45, ORF80 and ORF107) had clearly higher levels of expression compared to the other ORFs in both families (Figure 5, S3). In the LS family, these genes reached their highest expression level at 18 hpi, before declining at 24 hpi. By contrast, in the HS family, their expression increased strongly between 18 and 24 hpi, showing a 4-to 9-fold increase and reaching levels up to 10 times higher than in the LS family at 24 hpi (Figure 5). All five genes belong to cluster 3, except for ORF13 (protein containing a signal peptide domain, predicted to be extracellular), which belongs to cluster 2 (Table S1).

**Figure 5:**
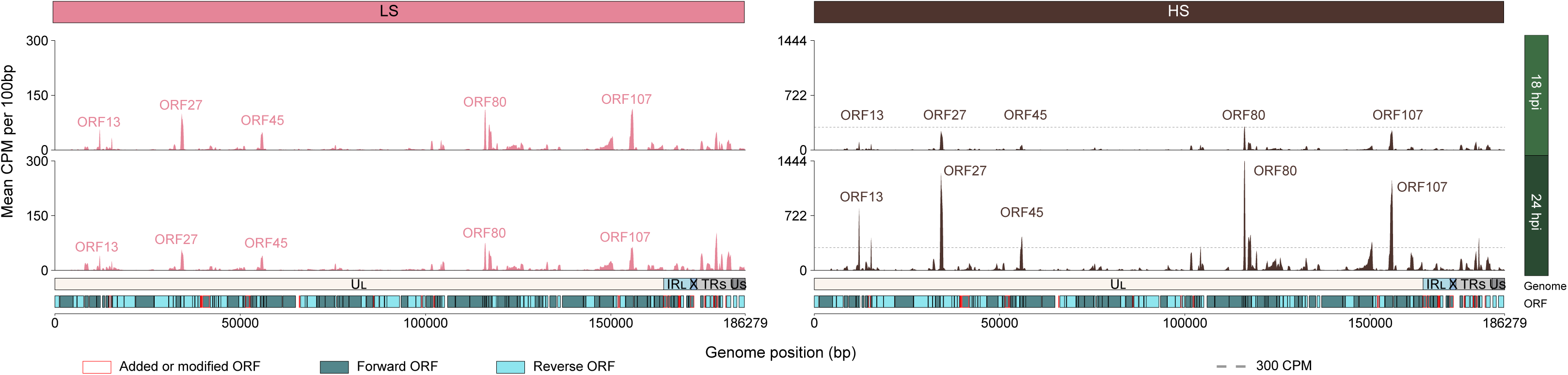
OsHV-1 whole genome expression over time and families. The mean Counts Per Million (CPM) per 100 bp windows is represented on the Y-axis for the 18 hpi library on the top and the 24 hpi on the middle. The plot of the LS family data, on the right, are colored in pink whereas the plot of the HS family data, on the left, are colored in darkbrown. The X-axis correspond to the OsHV-1 genome position in base paired. Representations of the OsHV-1 genomic regions (U_L_ in white, IR_L_ in light blue, X: in dark blue, IR_S_ in light grey and U_S_ in dark grey) and the ORFs location on the genome are shown above the X-axis. ORFs colored in darkblue are forward directed ORFs and ORFs colored in lightblue correspond to reverse directed ORFs. Finally, ORFs surrounding by red color correspond to new or modified ORFs compared to the OsHV-1 µVar A genome. The Y-axis of the LS family ranged from 0 to 300 CPM while it ranged from 0 to 1,444 CPM for the HS family. Dashed lines on plots of the HS family represent the maximum of CPM in the LS family.

#### OsHV-1 antisense/sense gene expression ratios differ between both Pacific oyster families

To investigate the putative regulation of gene expression by antisense transcripts, we analyzed the antisense/sense expression ratios of viral genes over time. Overall antisense/sense ratios remained stable across time (Figure S5). However, at later infection stages, some ORFs showed more sense than antisense reads in the LS family whereas the same ORFs showed more antisense than sense reads in the HS family. This is the case for ORF85 which had a ratio of 0.87 in the LS family and a ratio of 1.51 in the HS family at 18 hpi and 24 hpi. Similarly, ORF1, ORF9, ORF11, ORF22, ORFS32.1, ORF38, ORFS72.1, and ORFS115 showed a ratio from 0.59 to 0.97 in the LS family and a ratio from 1.18 to 3.05 in the HS family at 24 hpi (Figures 6, S3). With the exception of ORFS115, which has been identified as an origin-binding replication protein (Burioli et al., 2017), none of these ORFs have a predicted function. However, ORF1, ORF9, ORF85, ORFS72.1, ORF22, ORFS115 and ORF11 are predicted to localize to the nucleus, ORF38 to the cytoplasm and ORFS32.1 to the endoplasmic reticulum. ORF9 and ORF38 contain a zinc/RING finger domain, whereas ORFS32.1, ORF22 and ORF38 have a transmembrane domain. ORF22 and ORF11 contain a consensus disorder region and ORF11 also contains a coil domain (Table S1).

**Figure 6:**
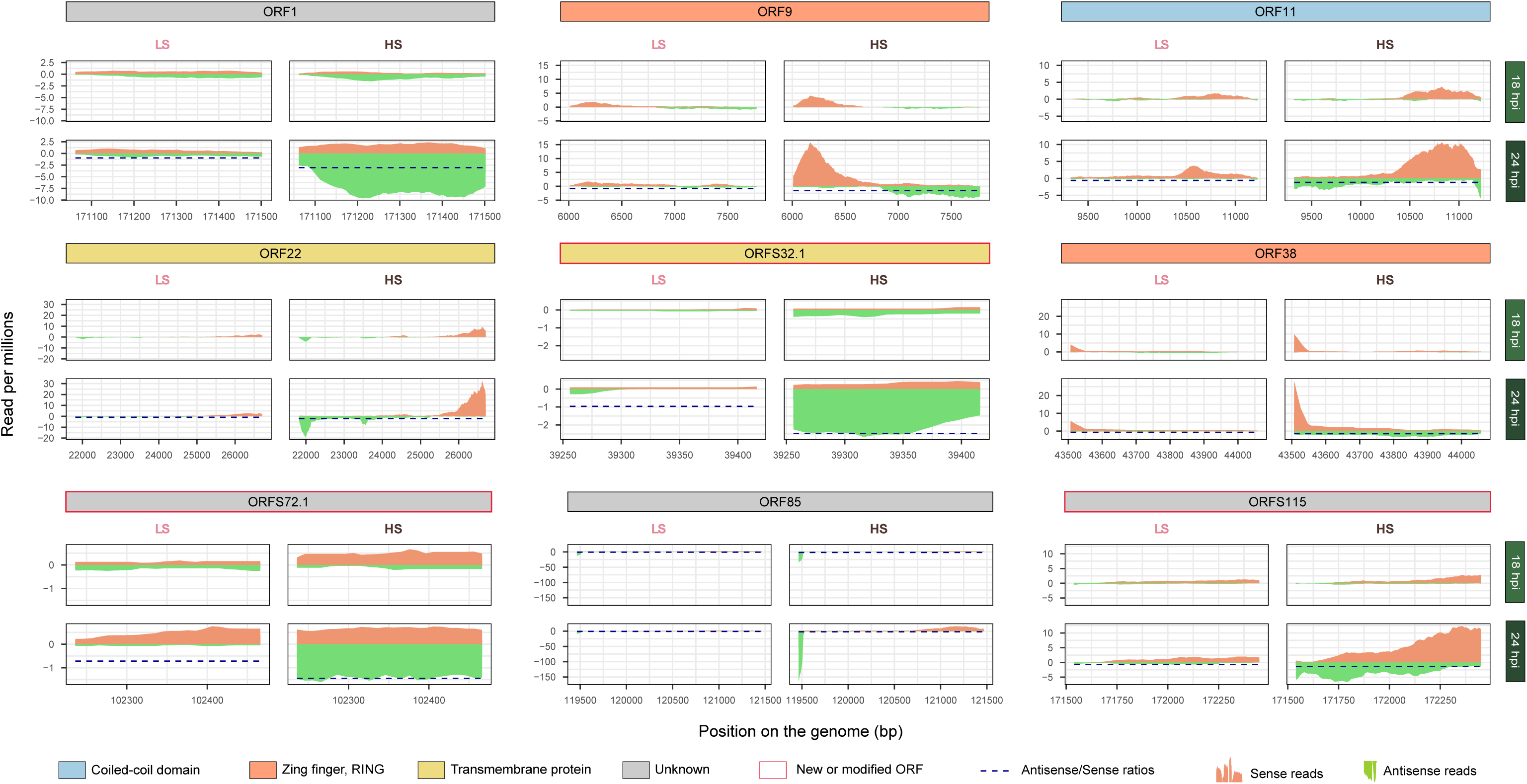
Sense and Antisense coverage for selected ORFs. Each panel represents a different ORF. The color of the header represents the protein predicted function or domain of each ORF. Red surrounding of the header means that the ORFs was new or modified compared to the OsHV-1 µVar A genome. For each panel the two graphs on the right represent the data for the LS family at 18 and 24 hpi whereas the two graphs on the left represent the data for the HS family at 18 and 24 hpi. The Y-axis of each graph represents the read count per millions and the X-axis represents the position of the ORF on the OsHV-1 genome. Orange coverage on the top of each graph represents the sense coverage while the green one represents the antisense coverage.

Notably, none of the ORFs showed the reverse pattern of having more antisense than sense reads in the LS family compared to the HS family.

#### OsHV-1 differential expression analysis reveals differences in infection dynamics between both Pacific oyster families

Statistical analyses were conducted to compare normalized expression levels between both families at each time point. During the early stage of infection (before 1 hpi), four genes were significantly overexpressed in the LS family: ORF49, ORF74 and ORF122 at 0.5 hpi and ORF49 and ORF90 at 1 hpi (Figure 7, Table S1).

**Figure 7:**
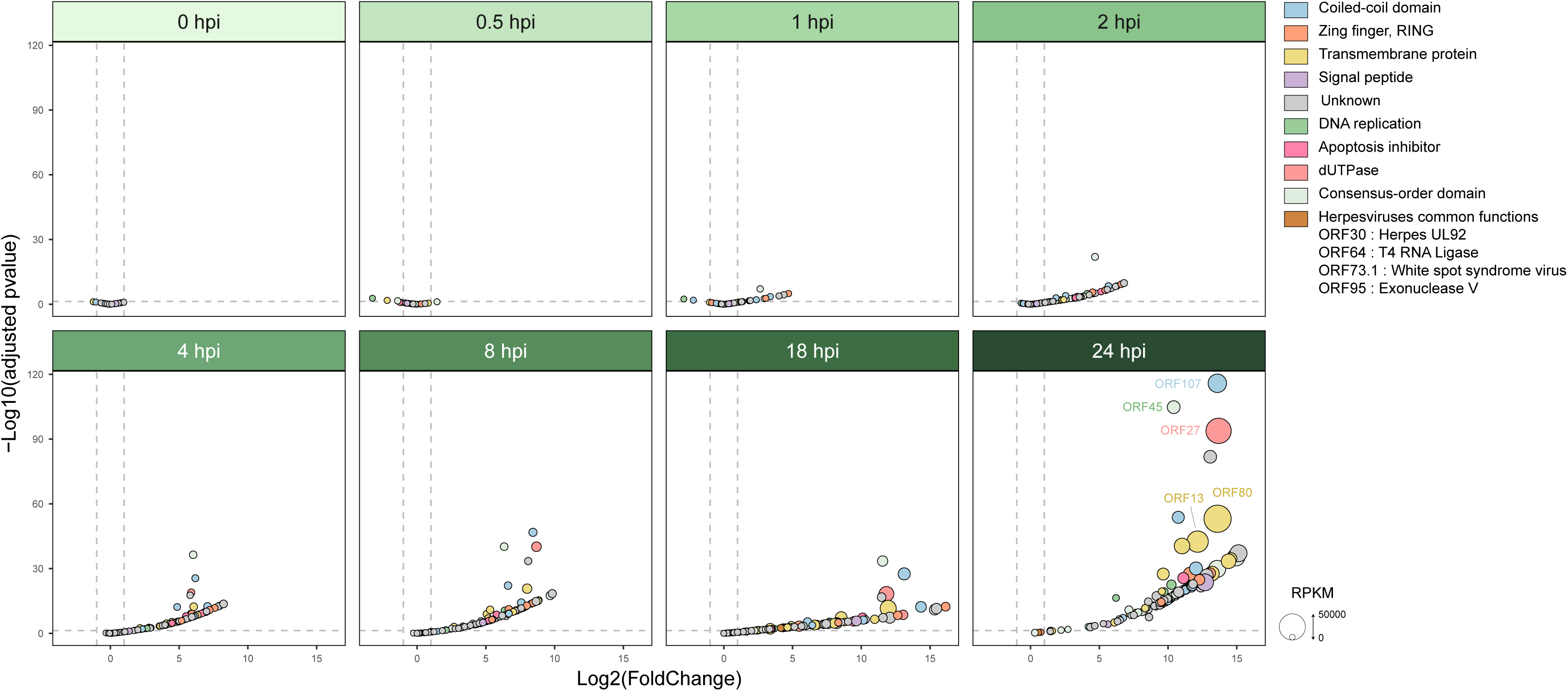
Volcano plot of differentially expressed viral genes. Differential expression analysis was performed using DESeq2 to compare viral gene expression between the HS and LS oyster families. The X-axis shows the log2 Fold-Change (Log2FC) and the Y-axis shows the negative log10 of the adjusted p-value (-Log10(padj)). The vertical dashed lines indicate a Log2FC threshold of ±1 and the horizontal dashed line marks the significance threshold for the adjusted pvalue padj = 0.05. Each point represents a viral gene, colored according to its function or domain. Point size reflects normalized expression in reads per millions per kilobases (RPKM). Genes to the left of the thresholds are overexpressed in the LS family, while those to the right are overexpressed in the HS family.

Then from 2 to 24 hpi all genes were overexpressed in the HS family compared to the LS one (Figure 7). Notably, genes coding dUTPase-like, ORF75 and ORF27, started to be overexpressed in the HS family at 1 hpi and 4 hpi respectively. Moreover, genes involved in the apoptosis inhibition (ORF42, ORF87, ORF99, ORF106) started to be overexpressed in the HS family at 2 hpi. Finally, for the genes involved in the DNA replication, ORF20, ORF67, ORF44 started to be overexpressed in the HS family at 2 hpi, ORF7, ORF66, ORF100 at 4 hpi, ORF109 at 8 hpi, 18 hpi and ORF49 at 24 hpi (Figures 7, Table S1).

### 3. Oyster genes expression

#### Haemocytes of both families respond to viral infection

To investigate Pacific oyster gene expression in response to OsHV-1 infection, we conducted a WGCNA analysis on infected libraries form both families. This approach identifies 59 genes modules, four of which (darkturquoise, grey60, honeydew and red) were significantly correlated with time and viral gene clusters expression (Figure S5).

The dark turquoise module, containing 5 057 genes, exhibited a robust negative correlation with viral clusters and time in the HS family, and a moderate negative correlation with time in the LS family (Figure S5). Gene Ontology (GO) enrichment analysis revealed that this module is primarily associated with cell mobility and enzymatic signaling, encompassing processes such as locomotion, cell adhesion, and morphogenesis (Table S2). The value of the eigengene steadily declined over time in the HS family whereas in the LS one, it initially declined, reaching a minimum at 18 hpi, before rising by 24 hpi (Figure 8).

**Figure 8:**
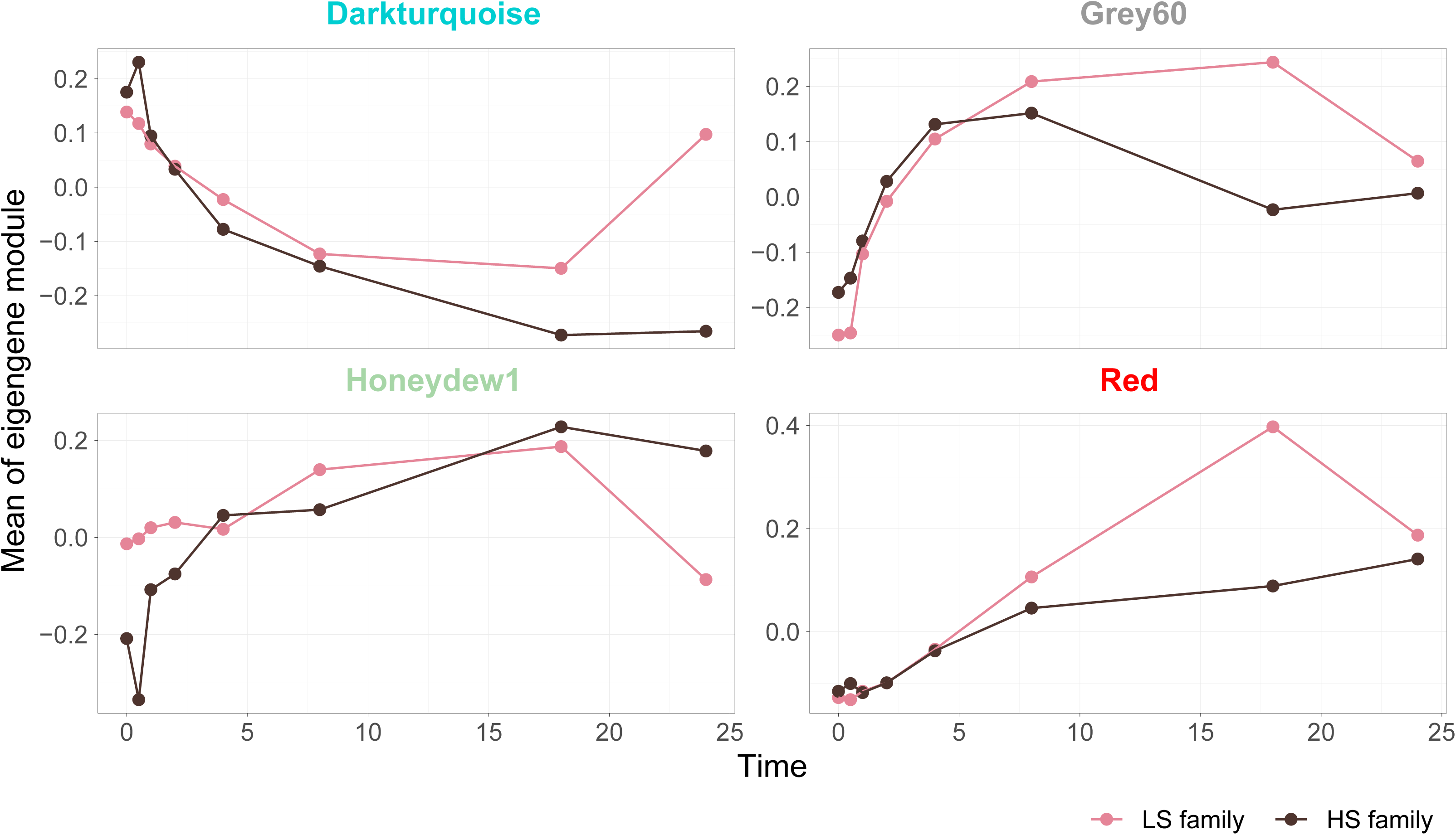
Eigengene value over time according to immune-related WGCNA modules. Each panel shows the eigengene profile of an immune-related module identified through WGCNA analysis. The Y-axis shows the mean eigengene value for each module and the X-axis shows time in hours post-infection (hpi). Pink lines and dots represent the LS family, while dark brown lines and dots represent the HS family.

The grey60 module, containing 1 340 genes, was positively correlated with both viral clusters and time in the LS family, and, to a lesser extent with time in the HS one (Figure S5). Go terms associated with this module were related to vesicular trafficking and Golgi apparatus organization (Table S2). For the HS family, the eigengene value increased until 8 hpi, then decreased through 18 hpi, and increased again towards 24 hpi whereas in the LS, it increased until 18 hpi, before decreasing by 24 hpi (Figure 8).

The honeydew1 module, containing 514 genes, showed a strong positive correlation with viral clusters and time in the HS family, and a moderate positive correlation with time in the LS one (Figure S4). Enriched GO terms suggested involvement in metabolic regulation, protein degradation, mitochondrial function and immune-related signaling pathways (Table S2). For the HS family, the eigengene value increased continuously whereas in the LS family, it rose until 18 hpi then decreased by 24 hpi (Figure8).

Finally, the red module, containing 1 668 genes, exhibited the strongest positive correlation with both viral clusters and time in both families (Figure S5). GO enrichment analysis revealed a dominant involvement with immune responses and the regulation of cellular stress (Table S2). The eigengene value increased slightly over time in the HS family whereas in contrast, it increased rapidly to 18 hpi in the LS family, before declining by 24 hpi (Figure 8).

#### Expression of most Pacific oyster genes drifts over the course of the experiment

Finally, to evaluate the effect of the experiment duration on Pacific oyster gene expression, we performed a WGCNA analysis on all control libraries. We identified 57 modules, eight of which were significantly correlated to the time. The black, darkviolet, darkturquoise, yellow and yellowgreen modules, containing 4 647 genes, showed a strong positive correlation with the time, whereas the magenta, grey60 and skyblue modules, containing 4 813 genes, exhibited a strong negative correlation with time (Figure S6).

GO enrichment analysis revealed that the negatively correlated modules are enriched for terms associated with the downregulation of key signaling pathways and the modulation of cellular architecture and chromatin dynamics. Conversely, the positively correlated modules were enriched for GO terms supporting mitochondrial bioenergetics and macromolecular assembly, including processes such as energy production, protein folding, and RNA processing (Table S3).

Among those 9 460 genes, 5 853 were shared with the immune-related modules identified previously (https://doi.org/10.6084/m9.figshare.29505254).

## DISCUSSION

The study aimed to improve our understanding of the OsHV-1 lytic phase in haemocytes from two Pacific oyster families that exibit different levels of susceptibility to OsHV-1 infection. First, we found that both viral DNA and RNA were detected in both families, indicating that OsHV-1 can infect and replicate in haemocytes. A similar pattern was observed in *M. gigas* tissue during OsHV-1 infection, with higher levels of viral DNA and RNA for HS Pacific oysters (Dégremont, 2011; Segarra, Mauduit, et al., 2014).

Then, to better understand both the dynamics of OsHV-1 replication and the immune response of haemocytes over time for the both families with contrasting responses to OsHV-1 infection, we used time-series dual RNA sequencing.

The expression of 144 OsHV-1 genes was monitored over a 24-hour period. Among them, 5 corresponded to modified ORFs from previous a previous OsHV-1 annotation (Burioli et al., 2017) and 17 were newly predicted ORFs. Nineteen of these 22 genes were expressed during the course of infection suggesting active transcription and a potential role in the lytic phase. However, three genes were not expressed suggesting that these genes were not expressed under these experimental conditions or because of incorrect ORF prediction.

To decipher the timing of the whole OsHV-1 gene expression, we first studied the viral replication over time. Clustering the viral genes into three groups, based on their expression, revealed that some viral genes were already expressed at 0 hpi in both families. Although this time point is slightly delayed due to the centrifugation step between the collection of haemocytes and RNA extraction, the detected gene expression suggests that OsHV-1 replication begins rapidly after infection. This observation indicates a faster onset of viral expression that previously described in haemocytes (1 hpi) or in the mantle (2 hpi) (Morga et al., 2017; Segarra, Mauduit, et al., 2014).

During the lytic phase of vertebrate herpesviruses in which this phase has been studied, gene expression follows a sequential pattern. Immediate-early genes are expressed shortly after the onset of infection and transcribed without the need of prior viral protein synthesis, then early genes are expressed at an intermediate stage and are transcribed with the help of immediate-early proteins and finally late genes are expressed at the end of infection once the replication of viral genome starts (Everett, 2014; Honess & Roizman, 1974; Knipe et al., 2013). Although these three categories may overlap in expression (Everett, 2014; Griffin et al., 2001) immediate-early and early gene expression typically declines during the infection (Beurden et al., 2013; Cherrington et al., 1991; Harkness et al., 2014; Honess & Roizman, 1974; Rozman et al., 2022; Tombácz et al., 2009).

Here, we identified 9 genes that are expressed immediately after in the onset of infection (i.e. after 0 hpi), 97 that are expressed early (i.e. after 1 hpi) and 38 that are expressed more lately (i.e. after 8 hpi). This represents an improvement on a previous study that analyzed the transcriptome of OsHV-1 using qPCR to target 39 ORFs, which did not allow classification into immediate-early, early, and late genes (Segarra, Faury, et al., 2014). In contrast, this whole-genome transcriptomic analysis of OsHV-1 provides a more precise evaluation of all viral ORFs, eliminating the bias introduced by targeting only a subset of genes via qPCR. Moreover, on the contrary to what have been shown for vertebrate herpesviruses, OsHV-1 gene expression increases over time and is cumulative. While this pattern could be specific to OsHV-1, it may be influenced by the presence of multiple haemocytes cell types in the primary culture that respond differently to viral infection, masking the sequential expression profile of early genes (Bachère et al., 1988; de la Ballina et al., 2022; Meng et al., 2022). For example, *M. gigas* granulocytes have been shown to be more immunocompetent to viral infection than hyalinocytes, which could lead to a delayed response to infection for this type of haemocyte within the overall cell population (Wang et al., 2017).

However, the expression of some OsHV-1 genes harbor similarities with vertebrate herpesviruses. In particular, genes coding for a dUTPase-like protein, apoptosis inhibitor and genes involved in the DNA replication belong to clusters that are characterized by an expression early after the onset of infection in Pacific oyster haemocytes, similarly to what have been observed in mantle of high susceptible Pacific oyster but also in human herpesviruses (Alharshawi et al., 2022; Knipe et al., 2013; Leopardi & Roizman, 1996; Segarra, Faury, et al., 2014; Wofford et al., 2020; Zarrouk et al., 2017). Interestingly, five genes harbor very high expression levels at the end of the infection with an even more pronounced trend in the HS family. Unfortunately, except for ORF27 that code for an dUTPase-like protein, none of the four remaining genes have a functional annotation. However, these genes contain predicted domain and predicted localization that suggest that they could be involved in the final assembly of the virion. For example, ORF80 possess a transmembrane domain and is predicted to be localized in the Golgi apparatus, which is known to be involved in the acquisition of the final envelope and tegument of the virion in herpesviruses and ORF13 contain a signal-peptide and is predicted to be located in the extracellular region and could be involved in the late stage of the progeny virion assembly (Crump, 2018; Everett, 2014; Knipe et al., 2013). Moreover, ORF27 was identified as highly expressed in mantle of another high and low susceptible Pacific oysters (Segarra, Mauduit, et al., 2014), proteins coded by ORF107, ORF27 and ORF45 have also been identified as highly present in another high susceptible Pacific oysters infected by OsHV-1 (Leprêtre et al., 2021) and ORF107, which is suggested to code for a capsid maturation protease, was among the most abundant gene expressed in OsHV-1 isolate ZK0118 infecting clam hemolymph (Rosani et al., 2024).

These findings suggest that these highly expressed genes may be involved in the assembly of the tegument and envelope of progeny virions released from infected cells. This is further reinforce by the observation that genes associated with virion assembly are among the most highly expressed in herpesviruses (Harkness et al., 2014; Knipe et al., 2013; Volkening et al., 2023). However, in other herpesviruses, genes involved in the assembly of the virion are generally classified as late genes (Beurden et al., 2013; Dotto-Maurel et al., 2025). In this study, four of these five genes were grouped into cluster 3, characterized by expression that occurs immediately after infection onset.

Stranded sequencing allowed us to investigate the potential role of antisense transcript in the regulation of viral expression. The analysis of antisense versus sense transcripts ratio shows that some genes harbor more antisense transcripts in the HS family than in the LS one at later stages of OsHV-1 infection. Antisense transcripts can interact with messenger RNAs (mRNAs) and may play a key role in post-transcription regulation. They may be involved in mRNA stabilization, regulation of translation, maturation, transport or localization and modification of proteins (Khorkova et al., 2015; X. Zhang et al., 2021). Antisense transcripts of lytic herpesvirus genes have been reported, although their function remains unclear. For example, approximately 50% of the viral genome of HSV-1 produce abundant symmetric transcripts at late stages of infection (Carter et al., 1996; Jacquemont & Roizman, 1975; Kozak & Roizman, 1975), while several antisense transcripts have been found overlapping ORFs in HCMV (G. Zhang et al., 2007). Here, the antisense versus sense ratios suggest that some OsHV-1 genes produce antisense transcripts. The observed differences between families may reflect a regulatory mechanism in the HS family aimed at slowing viral replication to prevent rapid cell destruction giving the virus a longer time frame to replicate before cell death.

To further understand the interactions between the virus and its host, we next investigated the transcriptional response of the pacific oyster to the OsHV-1 infection. By comparing the gene expression in haemocytes from both Pacific oyster families of infected libraries, we aimed to identify host pathways that are modulated during OsHV-1 infection and to determine how these might contribute to the contrasting levels of susceptibility. to viral infection. To this end, we characterized co-expression modules of host genes expression correlated with OsHV-1 infection for each contrasted oyster family, identifying four modules that highlighted distinct patterns depending on the family. The first module contained genes associated to the cell mobility, adhesion and morphogenesis, the second module included genes involved in vesicular trafficking and Golgi apparatus function and the final two were enriched in genes related to the immune response and the regulation of cellular stress.

On the first hand, in the LS family, the expression of genes involved in cell mobility, adhesion and morphogenesis decreased until 18 hpi before increasing sharply. In contrast, genes associated with vesicular trafficking, the Golgi apparatus, the immune response and cellular stress regulation showed a marked increase until 18 hpi followed by a decline. Furthermore, expression levels of the three viral gene clusters increased in the LS family until 18 hpi and then stabilized or decreased.

In the HS family, the expression of genes involved in cell mobility, adhesion and morphogenesis decreased until the end of experiment. In contrast, genes associated with vesicular trafficking, the Golgi apparatus, the immune response and cellular stress regulation increased until the end of experiment. Moreover, expression levels of the three viral gene clusters increased in the HS family until the end of experiment.

Taken together, these findings suggest that haemocytes for LS oysters activate a more effective and regulated immune response than HS oysters, thereby limiting the viral replication after 18 hpi. This response appears to be accompanied by a reduction in vesicular trafficking and Golgi activity, while processes related to cellular motility, adhesion, and morphogenesis are reactivated. In contrast, the immune response of haemocytes from the HS oysters may be too slow to control the infection as OsHV-1 genes are already highly expressed and viral particles are widely distributed within the haemocytes. These patterns have been previously observed with oysters from another LS family which responded earlier to the OsHV-1 infection (before 24 hpi), while oysters from another HS family responded later (after 24 hpi) (de Lorgeril et al., 2018).

Then, in order to evaluate how haemocytes change over the course of the experiment, we compared the Pacific oyster gene expression of control libraries from both families. We identified five gene modules positively correlated with the time and three negatively correlated. Gene Ontology enrichment analysis revealed that haemocytes initially mobilise resources to survive in culture, but gradually lose regulatory balance, resulting in progressive functional decline. As no viral material was detected in these control libraries, it is likely that the duration of the experiment affected the survival potential of haemocytes. Furthermore, 68% of the genes in the immune-related modules were also present in the time-related modules, confirming our redundancy analysis where the time accounts for 14.40% of the explained variance, whereas OsHV-1 infection explains only 8.91%. Taken together, these results imply that haemocytes are more affected by prolonged culture stress than by OsHV-1 itself and suggest that primary haemocyte cultures may not be suitable for extended time-series experiments.

Finally, for the first time, we conducted a time-series dual RNA sequencing on viral-infected haemocytes from two oyster families harboring different level of susceptibility to OsHV-1 infection. This study deciphers the sequential pattern of viral genes expression, providing new insights into the replication dynamics of OsHV-1 in Pacific oyster haemocytes. It also reveals how the immune response to infection varies between families, highlighting the role of host genetics in shaping the outcome of infection. However, our results also suggest that haemocytes maintained in artificial seawater may exhibit time-dependent stress responses that may confound infection studies. This highlights the importance of experimental conditions and suggests that future time-series analyses should take into consideration for cellular stress unrelated to viral infection.

## DATA AVAILABILITY

All sequence data have been deposited in ENA under the accession number PRJEB88219.

Analysis scripts are publicly available at https://gitlab.ifremer.fr/asim/haemocytes-dualrna-seq-analysis.git

## ACKNOWLEGDMENTS

### 4. Authors contribution

BM designed the experiment, LD provided the biological materials and NF and BM processed the samples. NF, BM and MT prepared the samples for sequencing and sent them. ADM, GC and JLL performed the bioinformatic analysis. ADM and GC drafted the manuscript, which was then corrected by all authors. All authors read and approved the final version of the manuscript.

### 6. Funding information

ADM and MT were financially supported by grant from the Ifremer Scientific Board. This work received financial support from the European projects VIVALDI (H2020 n°678589) and from the Ifremer’s Scientific Direction (Projet direction scientifique) and the EU funded project. The funders had no role in study design, data collection and interpretation, or the decision to submit the work for publication.

## Acknowledgments

We thank the staff of the Ifremer station at La Tremblade (ASIM) with a special thanks to Sandy Picot and Claire Martenot for their help with haemolymph punctions and Frédéric Girardin and his team (Plateforme des Mollusques Marins de La Tremblade PMMLT) and Christophe Stavrakakis and his team (Plateforme des Mollusques Marins de Bouin PMMB). We also thank the SEBIMER team for maintaining bioinformatics tools and the Pôle de Calcul et de Données Marines (PCDM; https://wwz.ifremer.fr/en/Research-Technology/Research-Infrastructures/Digital-infrastructures/Computation-Centre) for providing DATARMOR computing and storage resources.

## APPENDICES

**Figure S1: General statistics of sequenced libraries**.

The header shows the oyster family, time point, treatment (with nI: Non-Infected and I: Infected) and corresponding replicates for each library. The bar plots represent the total number of reads in each library, while the dot plots show the percentage of reads that aligned to the *M. gigas* genome (darkest pink or brown dots) to the OsHV-1 genome (lightest pink or brown dots) and reads that did not align to either genome (medium pink or brown dots). The Y-axis of the dot plot shows the log10 transformed percentage of reads.

**Figure S2: Heatmap of the normalized expression levels of the top 5000 expressed genes.**

Each line represents a gene, colored from red for high expression to blue for low expression. The header indicates the oyster family, time point, and treatment for each library. Asterisks represent outlier libraries that were removed for further analysis.

**Figure S3: Coverage of the OsHV-1 over time and families**.

The log10 of mean RPM (reads per million) per 100 bp windows is represented on the Y-axis for all the time point library. The plot of the LS family data, on the right, are colored in pink whereas the plot of the HS family data, on the left, are colored in darkbrown. The X-axis correspond to the OsHV-1 genome position in base paired. Representations of the OsHV-1 genomic regions and the ORFs location on the genome are shown above the X-axis. ORFs colored in darkblue are forward directed ORFs and ORFs colored in lightblue correspond to reverse directed ORFs. Finally, ORFs surrounding by red color correspond to new or modified ORFs compared to the OsHV-1 µVar A genome.

**Figure S4: Antisense/sense expression ratio for each viral gene in LS and HS family**.

The left panel shows the results for LS family and the right panel shows the results for the HS family. Each panel contains 144 facets representing the 144 ORFs following the genomic order. Header colors indicate genomic regions (white: U_L_ region, lightblue: IR_L_, darkblue: X, lightgrey: IR_S_, darkgrey: U_S_). In each facet, the X-axis shows the position on the OsHV-1 genome (in base pairs) and the Y-axis shows the antisense/sense expression ratio. Lines represent the ratio of antisense RPKM to sense RPKM, colored by time post infection. Circles indicate gene expression levels (sense and antisense) in RPKM. ORFs surrounded by a red rectangle are those that show differences in antisense versus sense ratios across families and that are represented in Figure 8.

**Figure S5: WGCNA modules description for infected libraries**.

The left panel shows the number of genes per module with the X-axis indicating the number of genes while the right panel represents the correlation between the modules and the traits (i.e. family, viral cluster 1 eigenvalue, viral cluster 2 eigenvalue, viral cluster 3 eigenvalue, LS family time and HS family time). Relationship between modules and traits are color coded from red for strong positive correlations to blue for strong negative correlations. For each module, the correlation coefficient and the associated p-value (in parentheses) are shown. The module names appear between the two panels. The bold, coloured names correspond to the immune-related modules that have been identified.

**Figure S6: WGCNA modules description for control libraries**.

The left panel shows the number of genes per module with the X-axis indicating the number of genes while the right panel represents the correlation between the modules and the traits (i.e. time and family). Relationship between modules and traits are color coded from red for strong positive correlations to blue for strong negative correlations. For each module, the correlation coefficient and the associated p-value (in parentheses) are shown. The module names appear between the two panels. The bolded names correspond to the time-related modules that have been identified.

**Table S1: Description of OsHV-1 ORFs from the reference genome and the genome used in this study, onset of expression, sense and antisense expression in RPKM, DESeq2 results, annotation and count matrices**.

ORFs in bold have been modified from the reference genome whereas ORFs in bold italic are new compared to the OsHV-1 µVar A genome.

**Table S2: GO term enrichment of WGCNA immune-related modules.**

**Table S3: GO term enrichment of WGCNA of time-related modules**

